# Intimate genetic relationships and fungicide resistance in multiple strains of human pathogenic fungus *Aspergillus fumigatus* isolated from a plant bulb

**DOI:** 10.1101/2021.05.23.445375

**Authors:** Hiroki Takahashi, Sayoko Oiki, Yoko Kusuya, Syun-ichi Urayama, Daisuke Hagiwara

**Affiliations:** Medical Mycology Research Center, Chiba University, 1-8-1 Inohana, Chuo-ku, Chiba 260-8673, Japan; Molecular Chirality Research Center, Chiba University, 1-33 Yayoi-cho, Inage-ku, Chiba 263-8522, Japan; Plant Molecular Science Center, Chiba University, 1-8-1 Inohana, Chuo-ku, Chiba 260-8675, Japan; Faculty of Life and Environmental Sciences, University of Tsukuba, 1-1-1 Tennodai, Tsukuba, Ibaraki 305-8577, Japan; Microbiology Research Center for Sustainability, University of Tsukuba, 1-1-1 Tennodai, Tsukuba, Ibaraki 305-8577, Japan

## Abstract

Fungal infections are increasingly dangerous because of environmentally-dispersed resistance to antifungal drugs. Azoles are commonly used antifungal drugs, but they are also used as fungicides in agriculture, which may enable enrichment of azole-resistant strains of the human pathogen *Aspergillus fumigatus* in the environment. Understanding of environmental dissemination and enrichment of genetic variation associated with azole resistance in *A. fumigatus* is required to suppress resistant strains. Here, we focused on eight strains of azole-resistant *A. fumigatus* isolated from a single tulip bulb for sale in Japan. This set includes strains with TR_34_/L98H/T289A/I364V/G448S and TR_46_/Y121F/T289A/S363P/I364V/G448S mutations in the *cyp51A* gene, which showed higher tolerance to several azoles than strains harboring TR_46_/Y121F/T289A mutation. The strains were typed by microsatellite typing, single nucleotide polymorphism profiles, and mitochondrial and nuclear genome analyses. The strains grouped differently using each typing method, suggesting historical genetic recombination among the strains. Our data also revealed that some strains isolated from the tulip bulb showed tolerance to other classes of fungicide, such as QoI and carbendazim, followed by related amino acid alterations in the target proteins. Considering spatial-temporal factors, plant bulbs are an excellent environmental niche for fungal strains to encounter partners, and to obtain and spread resistance-associated mutations.

## Introduction

Azoles are versatile compounds that show outstanding activity against a wide range of fungi, including plant and human pathogens. These compounds play an essential role in agricultural and clinical settings as fungicides and antifungal drugs (Fisher et al, 2018, Price et al, 2015). Their main mode of action is inhibition of the ergosterol biosynthesis pathway by inhibiting Cyp51, which functions as an 14-alpha-demethylase critical for the biosynthesis. Azole fungicides, known as demethylase inhibitors (DMIs), include triazole and imidazole compounds such as tebuconazole, propiconazole, triflumizole, and prochloraz. They are widely used to protect crops and fruits against pathogens by application during cultivation and postharvest preservation, as well as for seed disinfection. In medicine, azole drugs are essential options to combat dermatophytes and deep-seated fungal pathogens, such as *Trichophyton rubrum* and *Aspergillus fumigatus*, respectively. Azoles are the only class of compound used to control fungi in both agriculture and medicine.

*A. fumigatus* is a major causative agent of aspergillosis and ubiquitously present in the environment as a saprobe. A limited number of antifungals are approved for therapy of *A. fumigatus* infection; voriconazole (VRCZ) and itraconazole (ITCZ) are the first-line drugs for the treatment of pulmonary infection (Jenks and Hoenigl, 2018). However, this antifungal therapy is threatened by azole-resistant *A. fumigatus*, strains of which have been increasingly isolated since the beginning of this century (Howard et al, 2009). The resistance mechanisms to azole drugs that have been identified in *A. fumigatus* from clinical settings are mutations in Cyp51A, HMG-CoA reductase HMG1, and a subunit of CCAAT-binding complex HapE, and overexpression of *cdr1B*, which encodes an ABC transporter (Hagiwara et al, 2016a, Nywening et al, 2020, Hagiwara et al, 2018, Rybak et al, 2019, Camps et al, 2012, Hortschansky, et al, 2020, Fraczek et al, 2013). These azole resistance mutations are thought to have emerged during therapy with prolonged azole treatment.

However, in addition to treatment-based resistance, environmentally derived resistance has been considered as a non-negligible source of azole drug resistance of *A. fumigatus* during the last decade (Berger et al, 2017, Lestrade et al, 2019). Typical resistant strains from the environment carry a tandem repeat (TR) and single-nucleotide polymorphisms (SNPs) in the promoter and coding regions of the *cyp51A* gene, respectively. The most prevalent variants are TR_34_/L98H and TR_46_/Y121F/T289A, which were isolated for the first time from patients in Europe in 1998 and North America in 2008, respectively (Jeanvoine et al, 2020). The mutants with TR_34_ typically show high resistance to ITCZ, whereas the strains with TR_46_ show VRCZ resistance, but some are pan-azole-resistant strains. These genotypes were later recovered from environments worldwide (Schoustra et al, 2019, Resendiz et al, 2018, Hagiwara, 2018). Diverse resistant mutants with tandem repeats in the Cyp51A-encoding gene have been reported (Table 1).

**Table 1.**
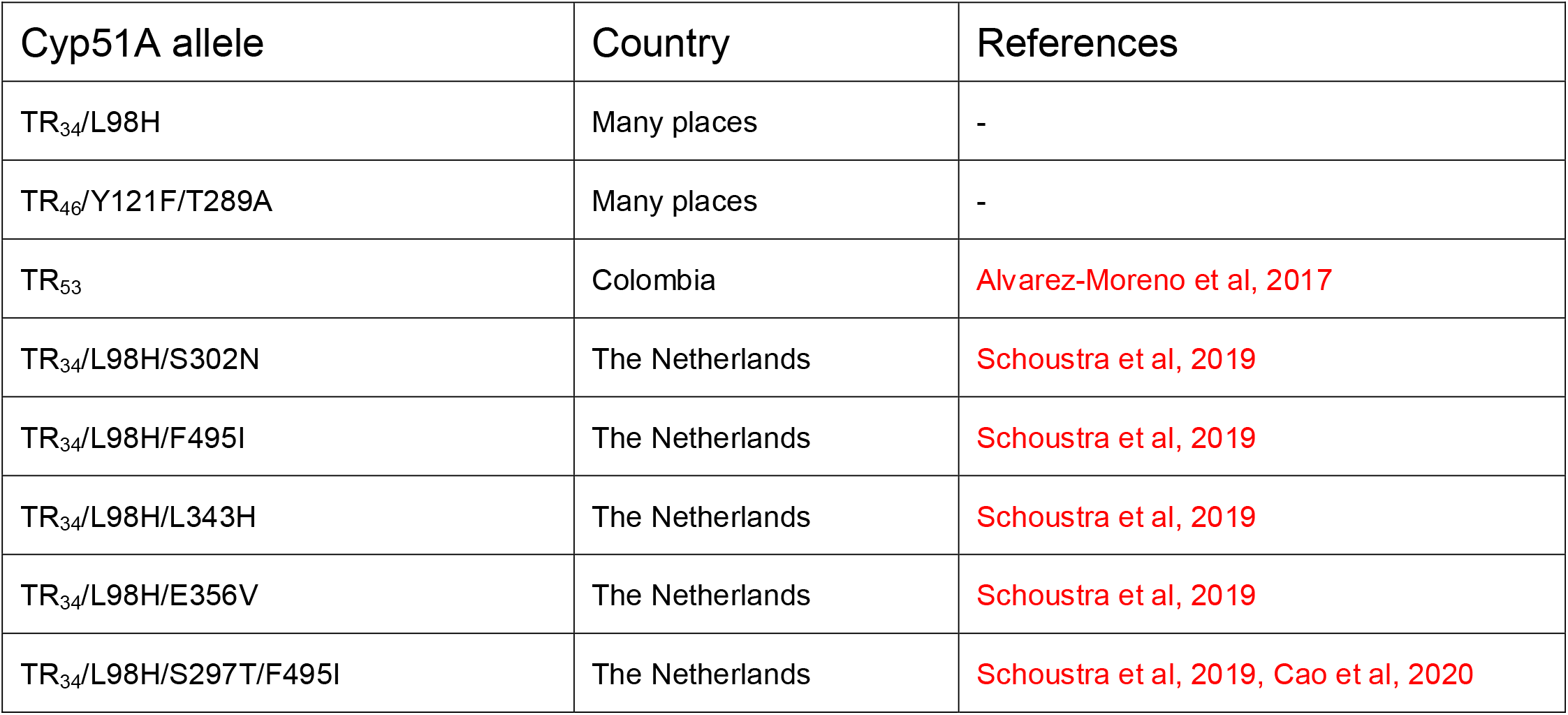
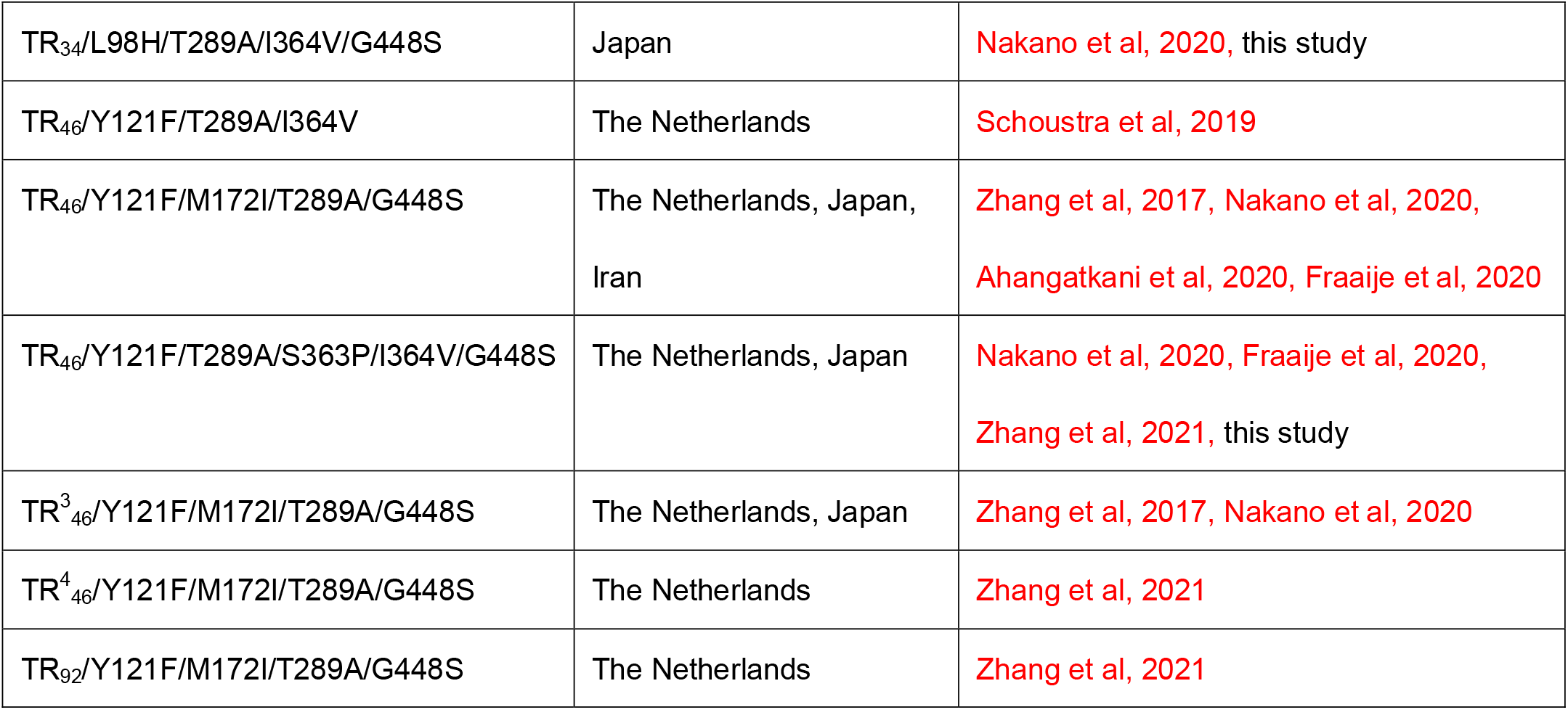
Reported Cyp51A variants with tandem repeats in azole-resistant *A. fumigatus*

Recently, a possible environmental hot spot for azole-resistant *A. fumigatus* was proposed (Zhang et al, 2017). The TR-type mutants were prevalently isolated from agricultural compost containing azole fungicide residues, whereas azole-free compost was dominated by wild-type (WT) *A. fumigatus*. This view was also supported in other studies (Schoustra et al, 2019, Zhang et al, 2021), indicating that azole-resistant strains are enriched under the selective pressure of environmental azoles. The work by Zhang *et al*. also suggested that sexual reproduction plays an important role in developing and evolving new *cyp51A* alleles for drug resistance in compost (Zhang et al, 2017). Taking into consideration that TR-type drug-resistant *A. fumigatus* mutants show cross-resistance to DMIs (Snelders et al., 2012), azole-containing environmental niches may serve as evolutionary incubators through genetic recombination.

The propagation of azole-resistant *A. fumigatus* has been studied in an epidemiological manner using microsatellite analysis by short tandem repeats for *A. fumigatus* (STR*Af*), which is a widely accepted intraspecies typing method with high-resolution discriminatory power (de Valk et al, 2005). TR-type mutant strains were spread worldwide. Some isolates from multiple countries were genetically closely related to each other and some had identical microsatellite patterns (Pontes et al, 2020, Cao et al, 2020, Wang et al, 2018, Hagiwara et al, 2016b). Besides such international propagation, intranational clonal expansion was also reported in several countries (Ahangarkani et al, 2020, Chowdhary et al, 2012). Recent population genomic studies revealed that the azole-resistant strains are globally distributed. The isolates were divided into two broad clades, and TR mutants belong to the populations in an uneven manner (Sewell et al, 2019). These data suggest that azole resistance primarily expanded by asexual and sexual propagation from a limited number of ancestors with TR-type mutation, rather than locally and independently emerging in each environment.

It was recently proposed that resistant *A. fumigatus* strains are transferred internationally via imported plant bulbs (Dunne et al, 2017). Plant bulbs produced in the Netherlands and sold in Ireland were contaminated with TR-type *A. fumigatus* mutants. Similar cases were also reported by two independent Japanese groups (Hagiwara 2020, Nakano et al, 2020); azole-resistant *A. fumigatus* with diverse Cyp51A variants were isolated from plant bulbs that were imported from the Netherlands and sold in Japanese gardening shops. These studies suggest that the wide spread of azole-resistant *A. fumigatus* mutants is attributable in part to trade in agricultural products including plant bulbs.

In the present study, to further understand genetic variations in plant bulb-associated isolates, we focused on eight *A. fumigatus* strains that were co-isolated from a single tulip bulb in a previous screening study (Hagiwara, 2020). Sensitivity to medical and agricultural azoles, as well as other classes of fungicides, was compared between the strains. Whole genome comparison of the eight strains showed several fragmental overlaps of their genomes, suggesting genetic recombination had occurred between strains in the single bulb. Our work indicates that plant bulbs are not only a vehicle for the pathogen but also a place where the pathogen can evolve its drug resistance.

## Results

### Variation of Cyp51A mutation in strains from a single bulb

In a previous study, eight strains of *A. fumigatus* were isolated from a single tulip bulb as different colonies (hereafter referred to as strains 3-1-A to 3-1-H) (Hagiwara, 2020). Strain 3-1-H has no TR or SNPs in *cyp51A*, whereas TR_34_ or TR_46_ occur in combination with various SNPs in the other seven strains (Table 2). Strains 3-1-A, 3-1-E, 3-1-F, and 3-1-G have a typical variant, TR_46_/Y121F/T289A. Strain 3-1-D has mutations S363P, I364V, and G448S as well as TR_46_/Y121F/T289A. Strains 3-1-B and 3-1-C have TR_34_/L98H and mutations T289A, I364V, and G448S. Notably, TR_34_/L98H and G448S are known to play a role in azole resistance, and T289A is typically accompanied by TR_46_ (Hagiwara et al, 2016a). Thus, the Cyp51A of strains 3-1-B and 3-1-C showed complicated sequence variation, including three mutations related to azole resistance.

**Table 2.** Cyp51A variation and microsatellite typing of the strains in this study

| Strain ID | Cyp51A variation | 2A | 2B | 2C | 3A | 3B | 3C | 4A | 4B | 4C |
| --- | --- | --- | --- | --- | --- | --- | --- | --- | --- | --- |
| 3-1-A | TR<sub>46</sub>/Y121F, T289A | 10 | 20 | 8 | 44 | 9 | 10 | 8 | 10 | 7 |
| 3-1-B | TR<sub>34</sub>/L98H, T289A, I364V, G448S | 23 | 10 | 9 | 35 | 9 | 6 | 8 | 10 | 18 |
| 3-1-C | TR<sub>34</sub>/L98H, T289A, I364V, G448S | 23 | 10 | 9 | 35 | 9 | 6 | 8 | 10 | 18 |
| 3-1-D | TR<sub>46</sub>/Y121F, T289A, S363P, I364V, G448S | 24 | 20 | 12 | 45 | 9 | 11 | 8 | 10 | 18 |
| 3-1-E | TR<sub>46</sub>/Y121F, T289A | 26 | 20 | 12 | 36 | 9 | 22 | 8 | 14 | 31 |
| 3-1-F | TR<sub>46</sub>/Y121F, T289A | 25 | 20 | 12 | 45 | 11 | 6 | 10 | 12 | 18 |
| 3-1-G | TR<sub>46</sub>/Y121F, T289A | 23 | 10 | 9 | 36 | 9 | 6 | 12 | 10 | 7 |
| 3-1-H | wt | 23 | 19 | 15 | 33 | 11 | 7 | 13 | 9 | 5 |

### Varied sensitivity to azoles in the strains from a single bulb

As previously reported, strains 3-1-A to G, which have TRs in *cyp51A*, showed VRCZ resistance (>32 µg/ml) in minimum inhibitory concentration tests (Hagiwara, 2020). To further understand the susceptibility to azole drugs, colony growth was evaluated on potato-dextrose-agar (PDA) containing 10 µg/ml of VRCZ (Fig. 1A). Strains 3-1-B, 3-1-C, and 3-1-D were more tolerant to VRCZ than the other strains. When grown on medium containing DMIs (triflumizole, imazalil, prochloraz, tebuconazole, epoxiconazole, or difenoconazole), strain 3-1-H, which harbors WT Cyp51A, showed the greatest growth inhibition among the strains. Strains 3-1-B, 3-1-C, and 3-1-D were less affected by the DMIs (except prochloraz) (Fig. 1B). On the basis of colony diameter measurement, strains 3-1-B, 3-1-C and 3-1-D showed higher tolerance to VRCZ and DMIs than strains 3-1-A, 3-1-E, 3-1-F, and 3-1-G (Fig. 1C). These results suggest that the combination of TR and G448S mutation increases resistance to azole compounds.

**Fig. 1.**
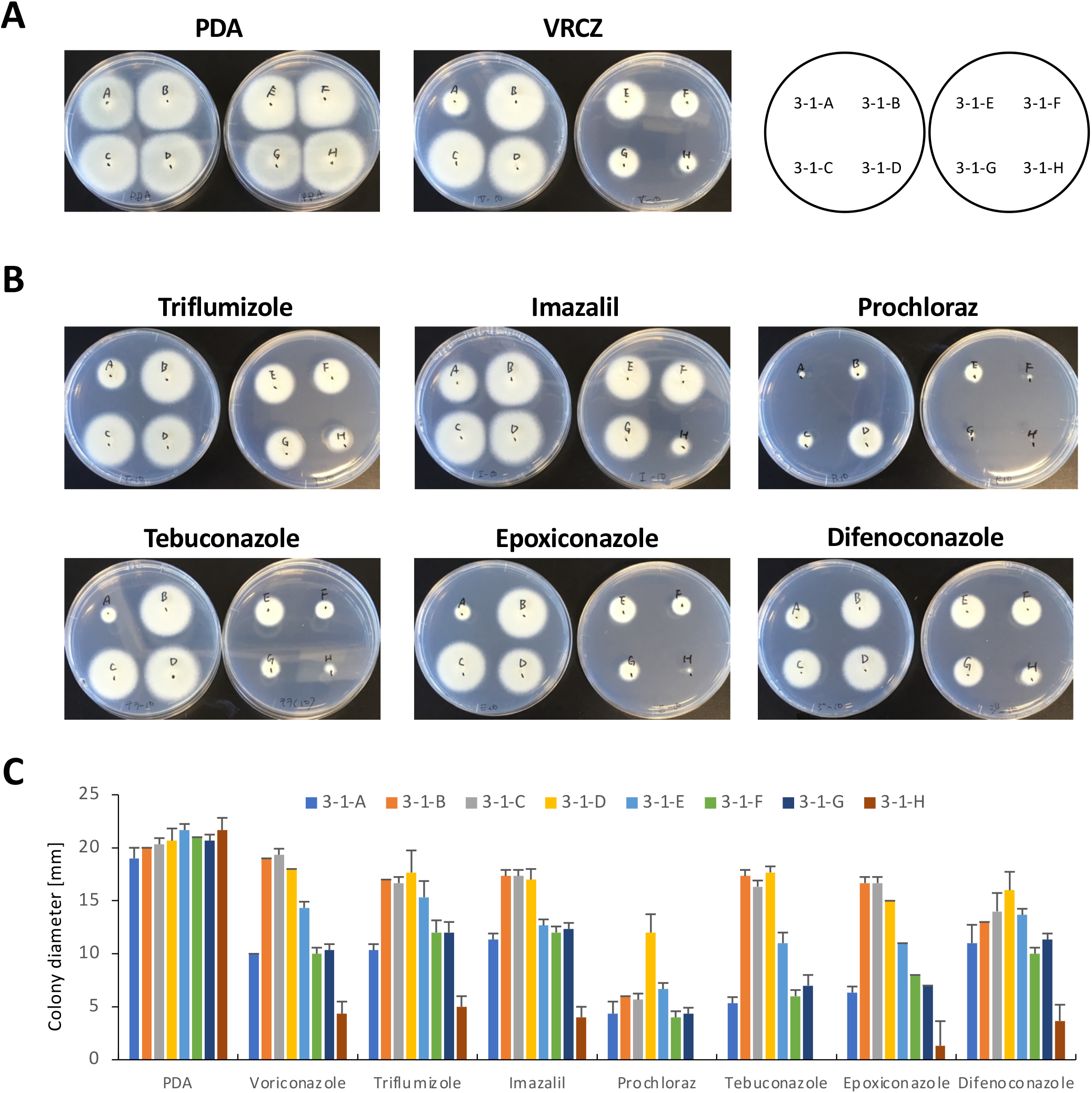
Colony growth of *Aspergillus fumigatus* strains isolated from a single tulip bulb on potato-dextrose-agar (PDA) containing azoles. (A) Growth on PDA containing voriconazole (VRCZ). Each strain was inoculated on PDA with dimethylsulfoxide (DMSO) as a control or 10 µg/ml VRCZ, and was incubated for 48 h. (B) Growth on PDA containing demethylase inhibitors (DMIs). Each strain was inoculated on PDA with DMSO as a control or 10 µg/ml DMI, and was incubated for 48 h. (C) Colony diameter on PDA containing VRCZ or DMIs. Each strain was inoculated on PDA with DMSO as a control or 10 µg/ml azole, and incubated for 28 h. Error bars represent standard deviations based on three independent replicates.

The expression levels of genes related to azole resistance were examined in the eight strains by quantitative real-time (qRT)-PCR. Compared with strain 3-1-H, which has the WT *cyp51A* gene, strains with a TR in the *cyp51A* gene showed higher expression of *cyp51A* (Fig. 2A). Overexpression of *cdr1B*, which encodes an ABC transporter, has been reported to confer azole resistance. Thus, the expression level of *cdr1B* was also determined in the eight strains by qRT-PCR. Strains 3-1-B and 3-1-C showed relatively high expression levels of *cdr1B* (Fig. 2B).

**Fig. 2.**
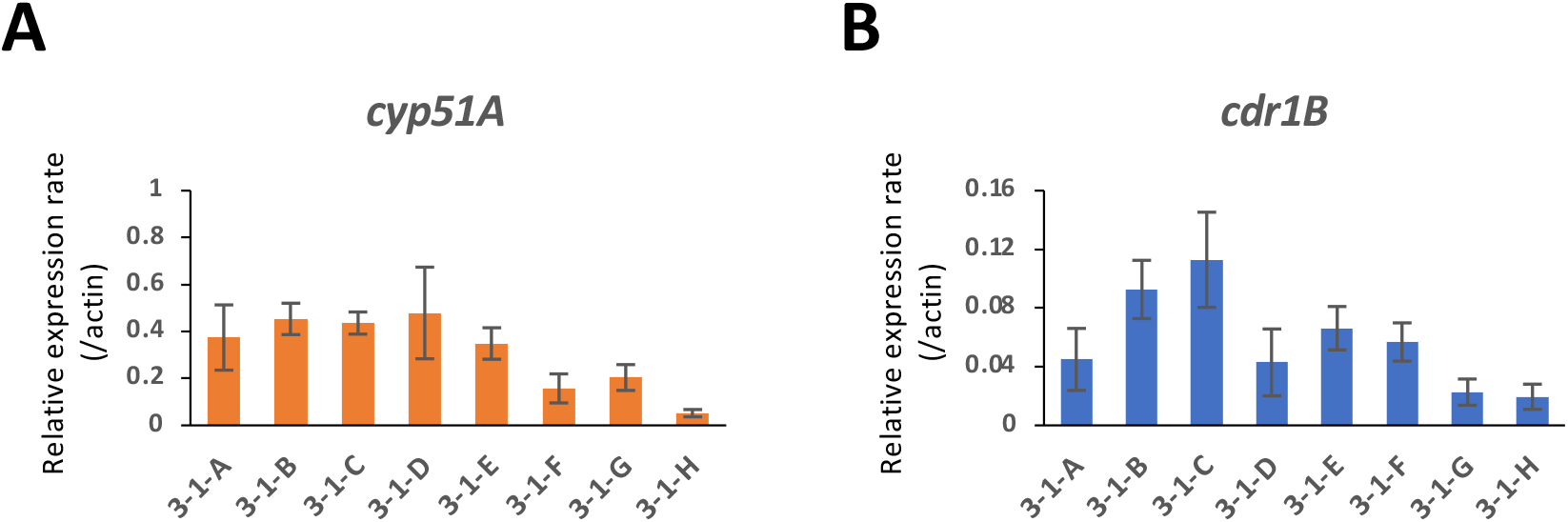
Gene expression analysis by quantitative real-time (qRT)-PCR. Expression levels of *cyp51A* (A) and *cdr1B* (B) were determined in the eight strains isolated from a single tulip bulb. The strains were cultured in potato-dextrose broth for 18 h. The *actin* gene was used as an internal control. Error bars represent standard deviations based on three independent replicates.

### Microsatellite typing analysis of tulip bulb isolates

To investigate the genetic relationships between the eight strains co-isolated from a single tulip bulb, microsatellite analysis using STR*Af* was performed (Table 2). This analysis also included TR-type strains that were previously reported and isolated in different countries and strains isolated from plant bulbs in Japan (Nakano et al, 2020, Hagiwara, 2020) (Fig. 3). Among the eight strains, the STR*Af* patterns of 3-1-B and 3-1-C matched perfectly. Strain 3-1-D is closely related to them, as this strain contains the same number of STRs in 4 of the 9 panels. Similarly, strain 3-1-D shares the same number of STRs as strain 3-1-F in 4 of the 9 panels. These four strains grouped into the same clade. The other strains were distantly positioned in the dendrogram. Interestingly, some strains that were isolated from plant bulbs in the study by Nakano *et al*. (2020) showed a close relationship with our strains. NGS-ER15 had an STR pattern similar to that of our strains 3-1-B and 3-1-C (5 of the 9 panells), which is consistent with these strains having the same Cyp51A allele (TR_34_/L98H/T289A/I364V/G448S). Strains NGS-ER6 and NGS-ER7 of Nakano *et al*. (2020) are closely related to strains 3-3-A and 3-3-B that were isolated from a single another tulip bulb in our previous study (Hagiwara, 2020). Note that these extraordinarily close relatives were isolated from plant bulbs in different laboratories.

**Fig. 3.**
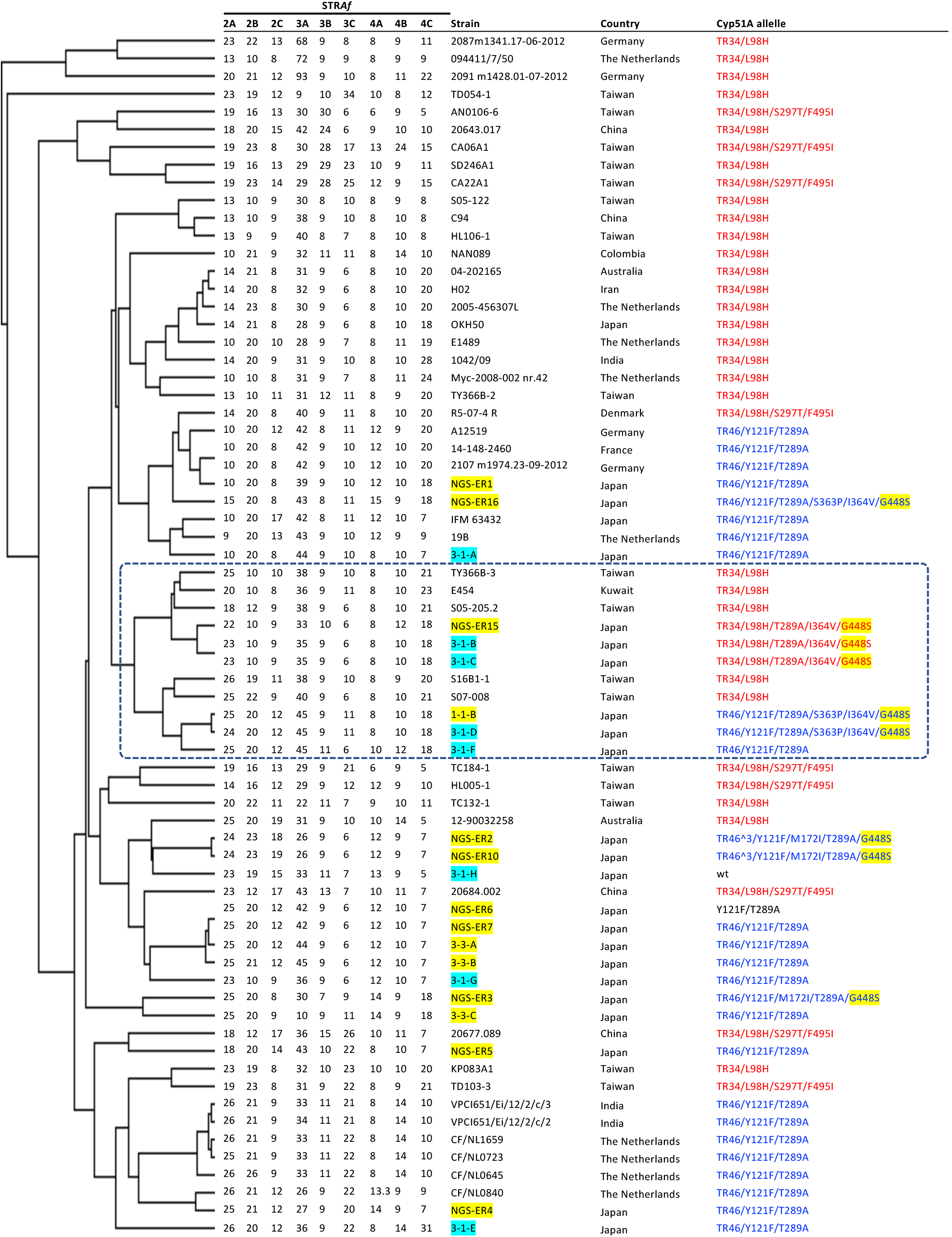
Microsatellite-typing analysis of *A. fumigatus* strains with tandem repeat (TR) mutations. The dendrogram was constructed using short tandem repeat for *A. fumigatus* (STR*Af*) patterns of the strains. The nine STR panels are shown. The strains listed refer to the literature (Chen et al, 2019, Hagiwara et al, 2016b, Nakano et al, 2020, Hagiwara, 2020). The names of strains isolated from plant bulbs in this study or other study are highlighted in pale blue or yellow, respectively. Cyp51A alleles with TR_34_ are indicated in red, and those with TR_46_ in blue.

### Genome sequencing and comparison between strains

To gain more insight into genetic differences or relatedness, genomes of the eight strains (3-1-A to H) were sequenced using the Illumina platform. Complete mitochondrial genomes were successfully obtained for the strains (31,749 to 31,770 base pairs [bp] long) (Table 3). A phylogenetic tree was constructed using the mitochondrial genomes and those of other strains (IFM 61407, IFM 59365, and IFM 61578) that had been clinically isolated in Japan (Takahashi-Nakaguchi et al, 2015) (Fig. 4A). This dendrogram indicated that the eight strains isolated from the tulip bulb can be divided into three groups. Group m1 contains strains 3-1-A, 3-1-D, and 3-1-G; strains 3-1-B, 3-1-C, 3-1-E, and 3-1-F are in Group m2. Strain 3-1-H was distantly positioned from both Group m1 and m2. Differences in the length of the mitochondrial genome well reflect the grouping, suggesting that strains within each group are very close relatives.

**Table 3.** Results of genome sequencing for strains isolated from a single tulip bulb

| Strain ID | Total length of chromosomes [bp] | GC [%] | # of proteins | Mitochondrial genome [bp] | Mito Group | CSP type | Mating type |
| --- | --- | --- | --- | --- | --- | --- | --- |
| 3-1-A | 28,889,155 | 49.342 | 9,492 | 31,770 | m1 | t02 | *mat1-1* |
| 3-1-B | 28,519,682 | 49.352 | 9,359 | 31,763 | m2 | t02 | *mat1-1* |
| 3-1-C | 28,533,261 | 49.355 | 9,490 | 31,763 | m2 | t02 | *mat1-1* |
| 3-1-D | 29,087,830 | 49.324 | 9,515 | 31,770 | m1 | t02 | *mat1-1* |
| 3-1-E | 28,703,796 | 49.398 | 9,464 | 31,763 | m2 | t02 | *mat1-1* |
| 3-1-F | 28,808,510 | 49.492 | 9,537 | 31,763 | m2 | t02 | *mat1-2* |

**Fig. 4.**
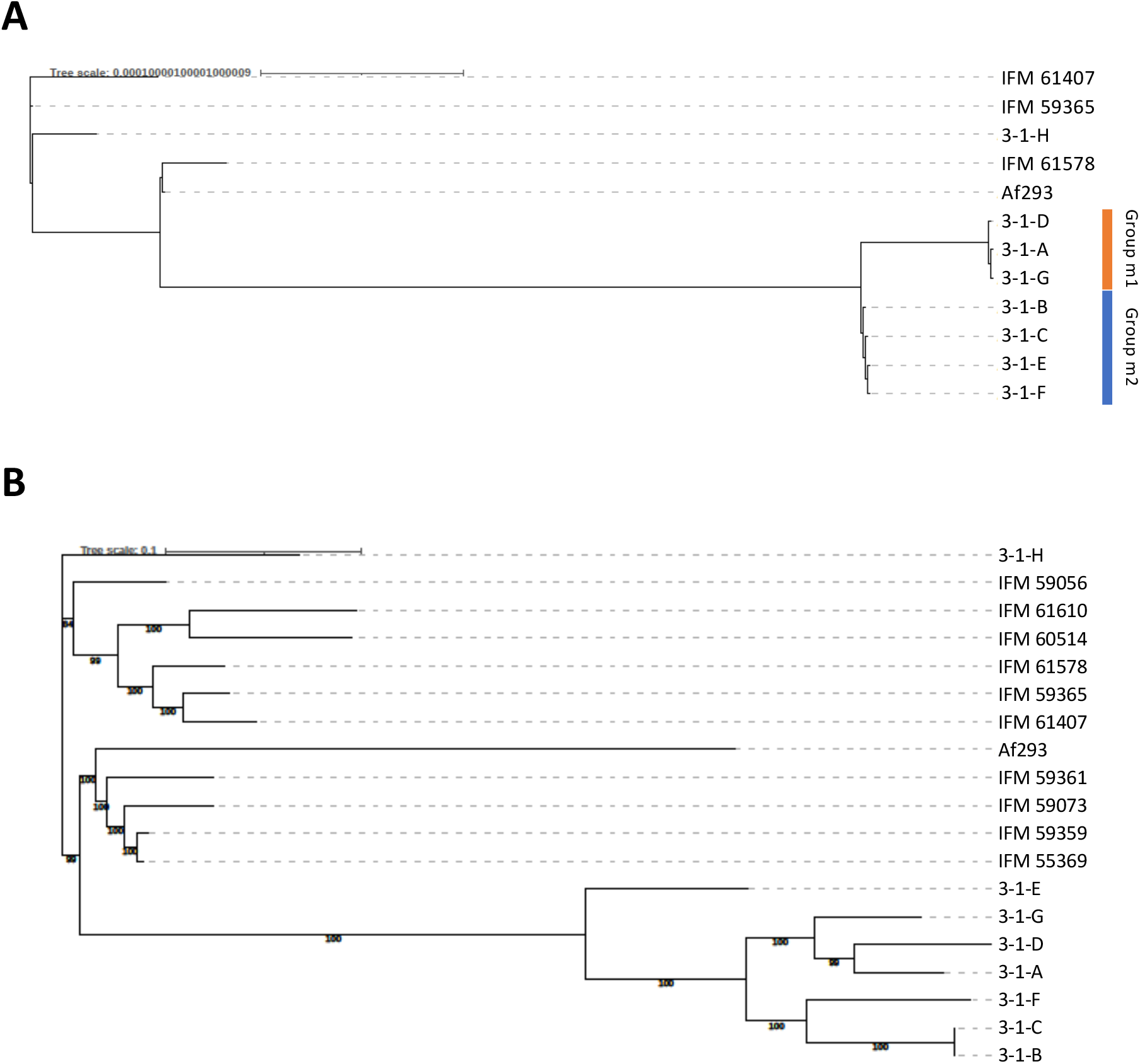
Phylogenetic trees constructed using mitochondrial (A) and nuclear (B) genomes. The trees were constructed using genomes of strains isolated from a single tulip bulb (3-1-A to 3-1-H) or clinically isolated in a previous study (Takahashi-Nakaguchi et al, 2016), as well as *A. fumigatus* reference strain Af293.

In the microsatellite typing analysis described above, strains 3-1-B, 3-1-C, 3-1-D, and 3-1-F were grouped into the same clade, but this was inconsistent with the grouping based on mitochondrial genomes, in which strain 3-1-D was not in the same group as strains 3-1-B, 3-1-C, and 3-1-F.

|  |  |  |  |  |  |  |  |
| --- | --- | --- | --- | --- | --- | --- | --- |
| 3-1-G | 29,178,518 | 49.295 | 9,543 | 31,770 | m1 | t02 | *mat1-1* |
| 3-1-H | 28,716,638 | 49.611 | 9,617 | 31,749 | m3 | t01 | *mat1-1* |

Nuclear genomes of the eight strains were compared with the reference genome of *A. fumigatus* strain Af293 (retrieved from AspGD, http://www.aspgd.org/); 92.2% to 93.6% of the Af293 genome was covered in the eight strains, and 69,949 to 79,391 SNPs were detected the genomes of the eight strains compared with the sequence of Af293 (Table 4). Phylogenetic analysis of the eight strains and previously-sequenced strains was performed by using concatenated sequences of the SNP positions (Takahashi-Nakaguchi et al, 2015) (Fig. 4B). Among the eight strains, 3-1-H was distantly positioned in the dendrogram as an independent clone. The other seven strains showed moderately close genetic-relatedness to each other based on comparison with the apparently independent clinical strains. Strains 3-1-B and 3-1-C showed the closest relationship, which was supported by the lowest number (6,241) of SNPs between strains (Table 4). This is consistent with the results of microsatellite and mitochondrial genome typing. Nevertheless, in the mitochondrial genome typing, strain 3-1-E was in Group m2 with strains 3-1-B, 3-1-C, and 3-1-F; however, strain 3-1-E was relatively distant from these three strains in phylogenetic analysis based on the nuclear genome (Fig. 4C).

**Table 4.**
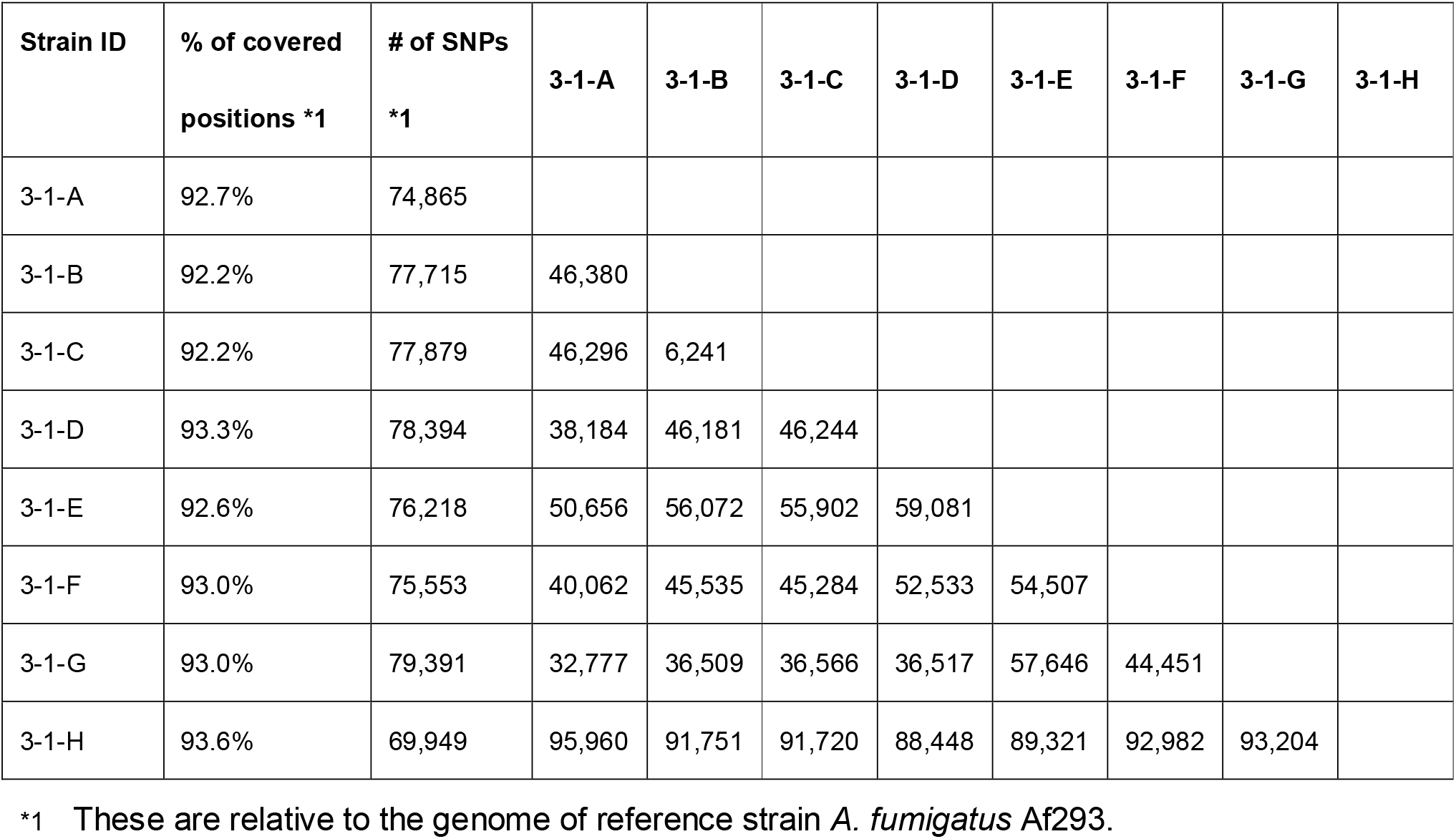
Summary of SNPs in strains isolated from a single tulip bulb.

From the genome sequences, CSP typing was performed, which can typify strains by sequence variation at a single locus (*csp*: Afu3g08990) (Klaassen et al, 2009). The results showed that seven strains (3-1-A to 3-1-G) carried an identical type (t02), but strain 3-1-H strain had type t01. Sequence analysis for mating type revealed that all but strain 3-1-F harbored *mat1-1*, whereas 3-1-F carried *mat1-2* (Table 3).

### Comparison of genome-wide SNP frequency pattern

Inconsistency in strain typing among the typing methods using the mitochondrial and chromosomal genome sequences caused us to speculate that genetic recombination had occurred between the strains isolated from the single tulip bulb. To help test this hypothesis, the SNP frequency and distribution were investigated and compared among the strains in a genome-wide manner (Fig. S1). There were several regions where the patterns of SNP frequency markedly differed among the strains (Fig. S1). For example, regions 5-A, 5-B, and 5-C on chromosome 5 were particularly characteristic (Fig. 5A). In region 5-A, strains 3-1-A, 3-1-E, and 3-1-H showed similar patterns of SNP frequency. In region 5-B, the pattern of strain 3-1-A was similar to that of strains 3-1-F and 3-1-H. In region 5-C, the pattern of strain 3-1-A was similar to that of strains 3-1-E, 3-1-G, and 3-1-H. These results indicate that strain 3-1-A shares parts of the sequence of chromosome 5 with strains 3-1-E, 3-1-F, 3-1-G, and 3-1-H. Such intergenomic variations were also found on other chromosomes (Fig. 5B, Fig. S1). These results showed a genome-wide mosaic pattern of SNP frequency, which is indicative of genetic recombination events in the strains.

**Fig. 5.**
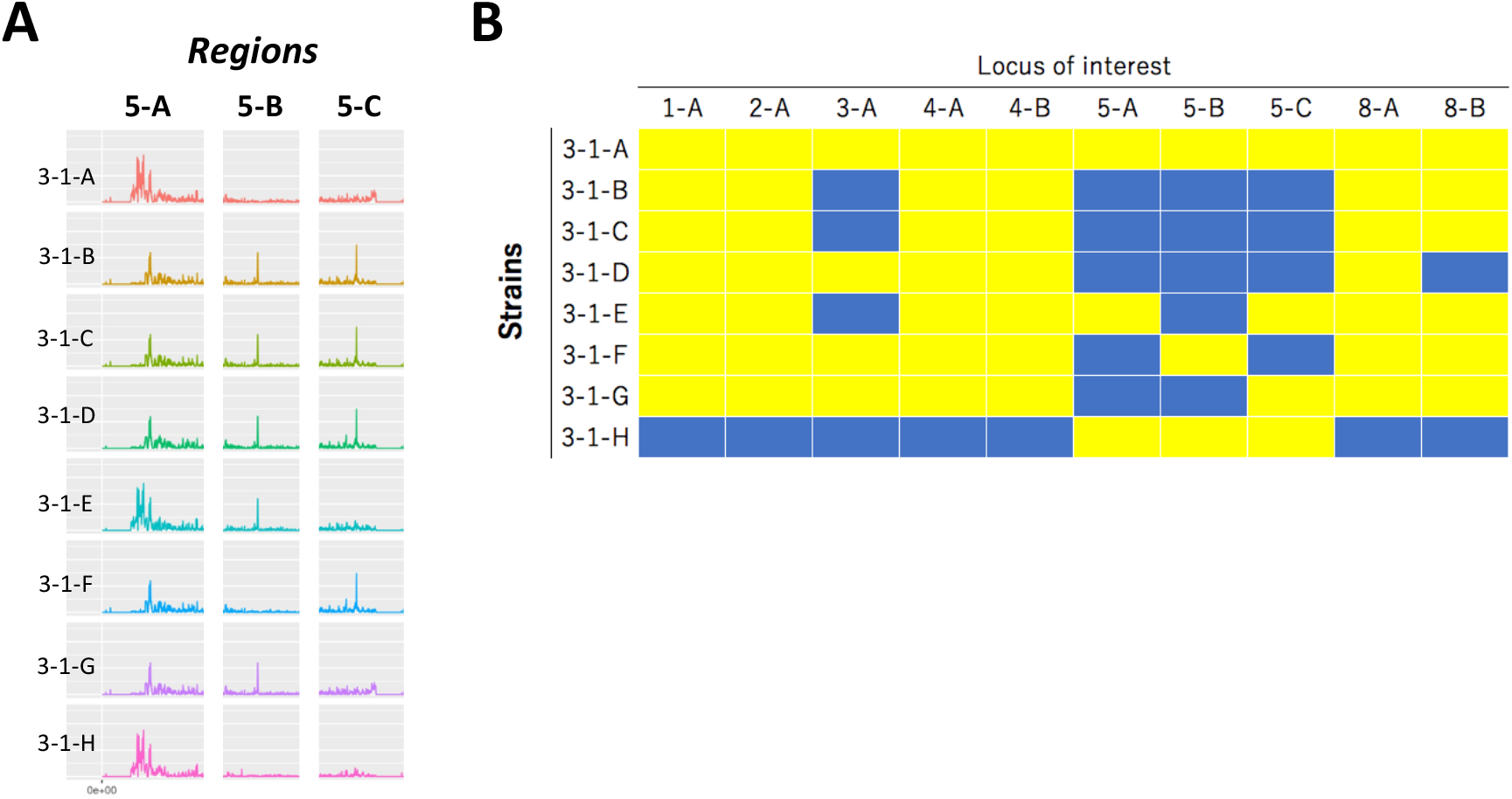
Differences in patterns of single nucleotide polymorphism (SNP) frequency among the strains isolated from a single tulip bulb. (A) The SNP presence patterns are compared in certain regions on chromosome 5 (5-A, 5-B, and 5-C). (B) The patterns in each region could typically be divided into two groups, which are indicated by yellow or blue panels. Ten genomic loci are shown and compared among the eight strains.

### Comparing genome-wide distribution of orthologous genes

To further investigate genome shuffling in the strains, we compared the patterns of orthologous among the strains isolated from the tulip bulb. First, the genes shared with the reference genome of *A. fumigatus* strain Af293 were investigated based on reciprocal blast hits (RBHs), which resulted in the isolates containing 8,196 to 8,322 orthologs of genes in strain Af293 (Table 5). The positions of the orthologs were generally evenly distributed in the strains, although fewer orthologs were found on chromosome 7. Notably, different patterns of ortholog content were displayed in some regions of the genomes of the various strains (Fig. 6). I.e., some sets of strains have lost particular sets of genes, and other sets of genes have been lost in other sets of strains. Hence, the set of strains that shares an ortholog pattern is different at each locus (Fig. 6). This suggests repeated genome shuffling among the strains.

**Table 5.** The number of orthologous genes in the strains isolated from a single tulip bulb compared with reference strain *A. fumigatus* Af293.

| Chromosome | # of orthologous genes compared with strain Af293 |
| --- | --- |

|  | Af293 | 3-1-A | 3-1-B | 3-1-C | 3-1-D | 3-1-E | 3-1-F | 3-1-G | 3-1-H |
| --- | --- | --- | --- | --- | --- | --- | --- | --- | --- |
| chr1 | 1,642 | 1,369 | 1,355 | 1,377 | 1,371 | 1,369 | 1,365 | 1,370 | 1,370 |
| chr2 | 1,640 | 1,433 | 1,406 | 1,425 | 1,425 | 1,411 | 1,436 | 1,427 | 1,433 |
| chr3 | 1,395 | 1,163 | 1,163 | 1,169 | 1,179 | 1,152 | 1,173 | 1,179 | 1,189 |
| chr4 | 1,253 | 1,089 | 1,075 | 1,093 | 1,090 | 1,098 | 1,096 | 1,085 | 1,095 |
| chr5 | 1,367 | 1,151 | 1,156 | 1,160 | 1,153 | 1,153 | 1,162 | 1,156 | 1,153 |
| chr6 | 1,249 | 1,068 | 1,067 | 1,074 | 1,058 | 1,069 | 1,073 | 1,070 | 1,080 |
| chr7 | 651 | 490 | 478 | 476 | 483 | 489 | 493 | 494 | 489 |
| chr8 | 628 | 502 | 496 | 507 | 497 | 505 | 506 | 491 | 513 |
| Total | 9,825 | 8,265 | 8,196 | 8,281 | 8,256 | 8,246 | 8,304 | 8,272 | 8,322 |

**Fig. 6.**
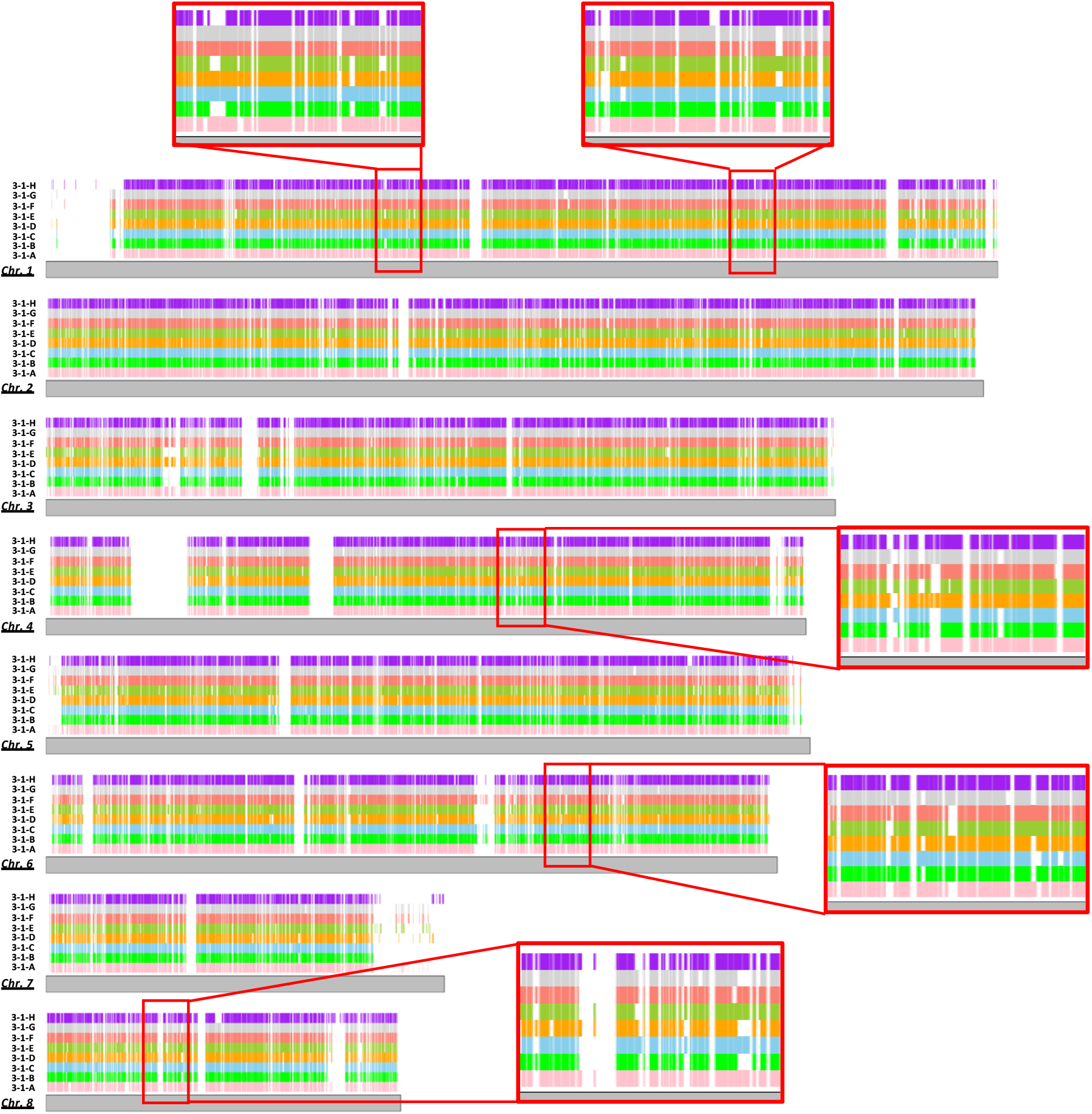
Visualization of genome-wide orthologous gene content in the strains isolated from a single tulip bulb. Orthologous genes were searched against the reference genome of *A. fumigatus* strain Af293. The presence of an ortholog is indicated by colored ribbons for each strain (3-1-A to 3-1-H). Some regions are enlarged to enable easier comparison of the patterns of gene content.

### Varied tolerance to agricultural fungicides

As these strains were derived from a horticultural product, they may have been exposed to agricultural fungicides besides DMIs. Hence, the susceptibility of the eight strains to QoI (pyraclostrobin), SDHI (boscalid), methyl benzimidazole carbamate (carbendazim), and phenylpyrrole (fludioxonil) was evaluated on PDA plates. There was no significant difference among the strains in susceptibility to fludioxonil and boscalid (Fig. 7A&B). However, the colony of strain 3-1-H was smaller than those of the other seven strains on the medium containing pyraclostrobin or carbendazim. These results suggest that there is varied tolerance to pyraclostrobin and carbendazim among the strains.

**Fig. 7.**
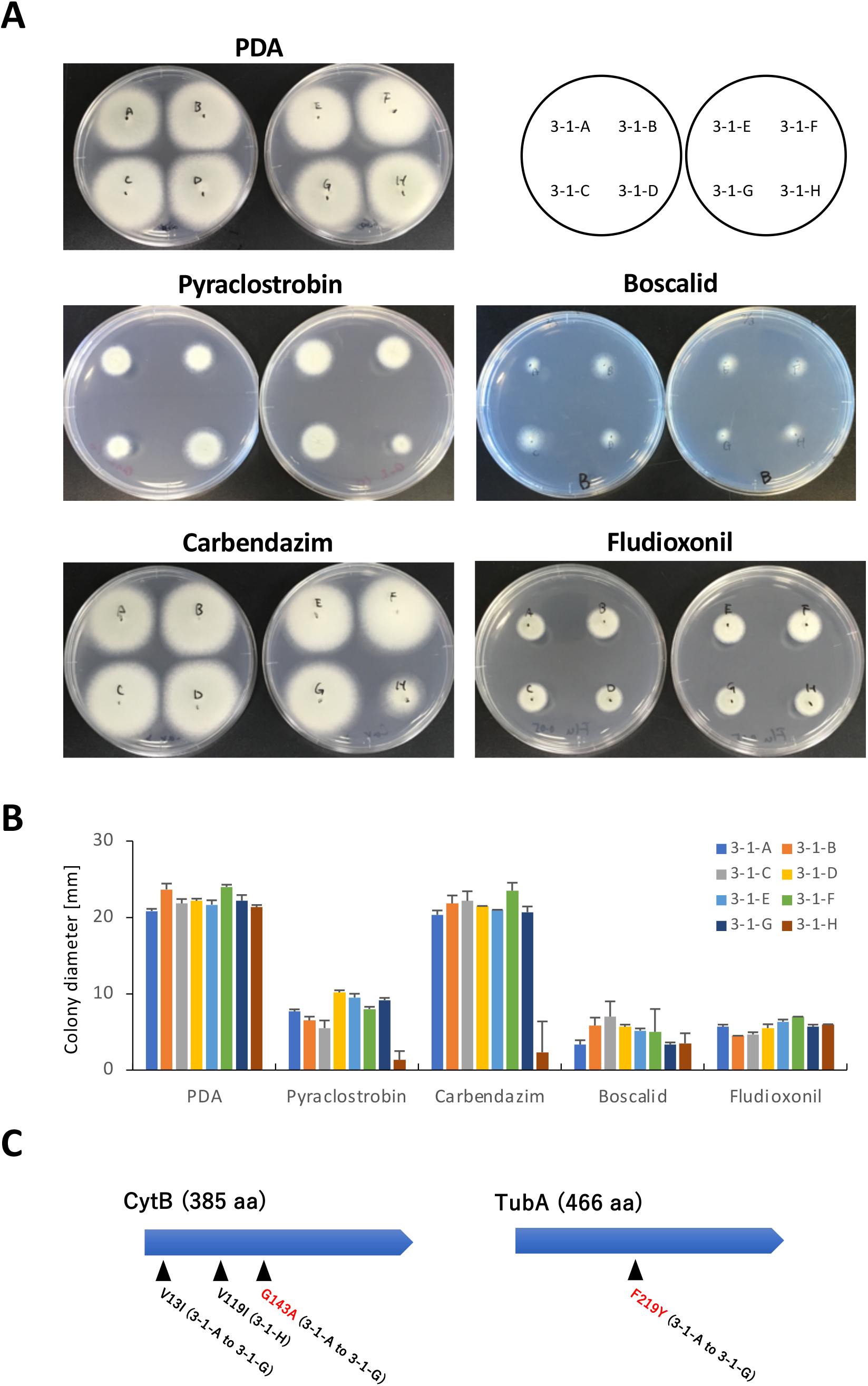
Colony growth of *A. fumigatus* isolated from a single tulip bulb on PDA containing fungicide. (A) Growth on PDA containing fungicide. Each strain was inoculated onto PDA with DMSO as a control, or QoI (pyraclostrobin; 10 μg/ml), SDHI (boscalid; 2.5 μg/ml), methyl benzimidazole carbamate (carbendazim; 5 μg/ml), or phenylpyrrole (fludioxonil; 0.2 μg/ml), and incubated for 48 h. (B) Colony diameter on PDA containing fungicide. Each strain was incubated for 30 h. Error bars represent standard deviations based on three independent replicates. (C) Amino acid substitutions detected in CytB and TubA of *A. fumigatus* strains isolated from a tulip bulb.

From the genome sequences, mutations that are possibly responsible for tolerance to carbendazim and pyraclostrobin were searched in the target molecules tubulin and cytochrome b that are encoded by *tubA* (Afu1g10910) and *cytB* (AfuMt00001), respectively (Fig. 6C). The amino acid substitution F219Y was found in TubA of strains 3-1-A to 3-1-G. This substitution has been reported in several carbendazim-resistant strains of plant pathogenic fungi (Yarden and Katan, 1993, Zhou et al, 2020). To investigate how the mutation is distributed in human pathogenic *A. fumigatus* genomes, the SNP database in FungiDB was explored. According to the dataset, 18% (14 of 77 strains) of *A. fumigatus* contain F219Y in TubA. Notably, eight of the 77 strains were isolated from the environment, and four of these possess the amino acid substitution.

In CytB, mutations V13I and G143A were found in strains 3-1-A to 3-1-G, and V119I was found in 3-1-H. G143A in CytB has been reported to confer the resistance to QoI in many plant pathogenic fungi (Samuel et al, 2011, Bolton et al, 2012), suggesting that this mutation in *A. fumigatus* is related to low sensitivity to QoI fungicide. As mitochondrial genome sequences are scarce in public databases, we investigated the sequence of the *cytb* gene in nine strains that were clinically isolated in a previous study and whose mitochondrial genome sequence is at least partly available (Takahashi-Nakaguchi et al, 2015). Among these nine strains, no G143A mutation was observed, whereas seven of the strains contain V119I (as also observed in our strain 3-1-H).

## Discussion

The distribution of azole-resistant *A. fumigatus* in natural environments has drawn increasing attention in recent years, with special interest in where the resistant strains have emerged, inhabit, and have been translocated to. However, deep understanding is still lacking. In this work, to fill in gaps in knowledge, we focused on strains that were isolated from a single tulip bulb.

Sexual reproduction of *A. fumigatus* was demonstrated in laboratory conditions in 2009 (O’Gorman et al., 2009). After this discovery, researchers paid more attention to the pan-genome of clinical isolates of this pathogenic fungus. However, gene flow in the environment has been poorly studied. Population genetics study using linkage disequilibrium analysis for genetic markers supported the view that *A. fumigatus* reproduces asexually and sexually in natural habitats (Klaassen et al., 2012). However, although there is accumulating evidence for genetic recombination in nature, proving the occurrence of sexual reproduction is difficult unless one can directly collect cleistothecia and ascospores formed in the environment. Genetic recombination in sexual development is suggested to cause the emergence of TR mutation in *cyp51A* gene through unequal crossover (Zhang et al., 2017). Thus, sexual reproduction is considered both to spread mutations by fusion with other strains and production of progeny, and to locally produce *de novo* TR mutations, which could affect the prevalence of resistance to drugs and fungicides. Our data show that seven of the eight isolates from a single tulip bulb contain TR mutations, and the genetic variation between the TR-containing strains is low compared with that between apparently independent strains. In addition, on the basis of genome-wide distributions of SNPs and orthologous genes, genetic recombination is likely to have occurred between the seven strains.

Co-isolation of the strains from a single bulb indicates that they have had much opportunity to physically interact with each other inside or on the bulb. In addition to the close spatial relationship of the fungal strains, they may interact for a long time. In the conventional process of plant bulb production, bulbs are multiplied from a parental bulb. This bulb multiplication is continued every year, which presumably causes the sustained presence of the fungi on/inside the bulbs. As described here, several strains were attached to a single bulb. These strains might have encountered others and genetically mixed many times. Once mutations giving rise to resistance to azoles emerged, the mutations could be preferentially and stably retained in the microbial community inside the bulb.

Sequencing analysis of the eight strains produced complete mitochondrial genomes and chromosomal genomes. The mitochondrial genome was of great help in interpreting whether there had been sexual reproduction among the strains. On the basis of mitochondrial genome sequences, the strains with TR mutation can be classed into Groups m1 and m2 (Fig. 4A). The length and sequences of the mitochondrial genomes are highly conserved in each group, indicating that they are genetically close progenies. However, the chromosomal genomes were diverse among the strains with TR mutation to the extent that there were 32,000 to 59,000 SNPs, excepting strains 3-1-B and 3-1-C which had approximately 6,200 SNPs (Table 4). The differences in grouping based on mitochondrial and chromosomal genomes strongly suggested a genetic recombination event. We therefore propose that the complete mitochondrial genome is valuable for gaining deeper insight into genetic relatedness among and between environmental and clinical isolates.

Strains with complicated *cyp51A* alleles have been reported in the literature and this paper (Table 1). For instance, we have isolated *A. fumigatus* strains with TR_34_/L98H/T289A/I364V/G448S (3-1-B and 3-1-C) and TR_46_/Y121F/T289A/S363P/I364V/G448S (3-1-D) mutations in the *cyp51A* gene. TR_34_/L98H is a typical TR-type mutation conferring resistance to ITCZ and in some cases to VRCZ, whereas TR_46_/Y121F/T289A confers resistance to VRCZ and in most cases to ITCZ (van Ingen et al, 2015, Buil et al, 2018). Amino acid substitution G448S contributes to resistance to VRCZ and occasionally to ITCZ (Bellete et al, 2010, Toyotome et al, 2016, Cao et al, 2020). Our finding that three strains (3-1-B, 3-1-C, and 3-1-D) showed a higher tolerance to VRCZ and some DMIs than strains with only TR_46_/Y121F/T289A mutation is suggestive of elevation of tolerance to azole drugs by combining mutations. Importantly, strains with G448S mutation have been isolated not only from clinical samples but also from soil (Cao et al, 2020). We cannot rule out the possibility that the G448S mutation originally emerged and was retained in strains with TR_46_/Y121F/T289A under the selective pressure of fungicides.

The *A. fumigatus* strains used in the present work were isolated from a tulip bulb by culturing at 45°C on plates containing medium supplemented with fluconazole to select fungi that were resistant to fluconazole (Hagiwara 2020). In total in that study, *A. fumigatus* was isolated from 50.8% of tulip bulbs (96/189), and strains isolated from 20.6% of the bulbs (39/189) had TR mutation. Because *A. fumigatus* is a saprophytic fungus that widely inhabits soil, compost, plant debris, wood chips, the air, and aquatic environments, it was not surprising that half of the tulip bulbs were contaminated with *A. fumigatus*. However, we have no idea how the fungus resides on or inside the bulbs from a biological viewpoint. Because of the high frequency of *A. fumigatus* isolation from tulip bulbs, there might be certain mechanisms by which *A. fumigatus* colonizes and infects the plant tissue, enabling persistence across bulb progenies. Notably, some *A. fumigatus* strains were isolated from *Citrus macrocarpa, Myricaria laxiflora, Ligusticum wallichii*, and *Moringa oleifera* (Francisco et al, 2020, Qin et al, 2019, Li et al, 2020, Arora and Kaur, 2019) as an endophyte. In general, however, the view that *A. fumigatus* has an endophytic mode in its life cycle remains to be established. In consideration of the dynamic mobilization of *A. fumigatus* in the environment, its association with plants may be overlooked, and we should pay more attention to it.

Recently, several field studies were published in which the prevalence of azole-resistant *A. fumigatus* was investigated in association with fungicide use. Work by Zhou *et al*. (2021) demonstrated that the concentration of triazoles in the soil of greenhouses was not significantly correlated with azole susceptibility of isolates. In another study from Germany, a low frequency of azole-resistant isolates from crop fields was reported regardless of azole fungicide use (Barber et al, 2020). A study by Frasije *et al*. (2020) also reported a low number of azole-resistant isolates in the soils of wheat-cropping fields subjected to fungicide treatment. The authors considered that arable crop production is low risk for development of azole resistance. Conversely, a large-scale survey across China was conducted, which showed that the residual level of azole fungicides in paddy soils positively correlated with the prevalence of azole-resistant *A. fumigatus* (Cao et al, 2021). Field research on azole-resistant *A. fumigatus* has started in many countries. More studies are required on the effects of fungicide use on the occurrence and spread of azole-resistant *A. fumigatus* in the environment, including agricultural and horticultural settings.

In plant bulbs, there may be other pathogenic and nonpathogenic fungi beside *A. fumigatus*. They are also exposed to fungicides when the bulbs are treated with fungicide. Repeated use of fungicides would facilitate the occurrence of resistance mutations in non-targeted fungi as well as in the target fungi of the pesticide. In the present study, we found that mutations in CytB and TubA that are related to resistance to QoI and carbendazim fungicides, respectively, were detected in *A. fumigatus* strains as an example of non-target fungi. These mutations might have been resulted from fungicide exposure during bulb production. Importantly, identical mutations of *A. fumigatus* were reported by Fraaije *et al*. (2020) and are found in database. These findings suggest that mutations related to resistance to antifungal agents are already present in the genomes of environmental fungi regardless of their pathogenicity. The boundary between acquired and natural resistance to antifungal compounds may become unclear in the near future.

## Materials and Methods

### Strains and culture conditions

Strains 3-1-A to 3-1-H used in this study were obtained in previous study and were isolated from a single tulip bulb (Hagiwara, 2020). For plate and liquid cultures, PDA and potato-dextrose broth (PDB) were used, respectively. For colony growth tests, 10^5^ conidia of each strain were inoculated and incubated for 48 h at 37°C before taking pictures. Insusceptibility tests, 10 µg/ml VRCZ, imazalil, prochloraz, triflumizole, tebuconazole, epoxiconazole, and difenoconazole were respectively added to PDA. The control plate contained the equivalent volume of dimethylsulfoxide (DMSO). For measuring colony diameter, the culture time was 28 or 30 h. The data were obtained in triplicate, and the mean and standard deviation are presented. The fungicides fludioxonil, carbendazim, boscalid, and pyraclostrobin were used at 0.2 µg/ml, 5 µg/ml, 2.5 µg/ml, and 10 µg/ml, respectively.

### Quantitative real-time RT-PCR

Strains were cultured in PDB at 37°C for 18 h and harvested. The mycelia were frozen in liquid nitrogen, and total RNA was isolated using Sepasol Super G (Nacalai Tesque, Kyoto, Japan). cDNA was obtained by reverse transcription reaction using the total RNA sample and ReverTra Ace qPCR RT Master Mix with gDNA remover (TOYOBO, Osaka, Japan).

Real-time RT-PCR was performed using Brilliant III Ultra-Fast SYBR Green QPCR Master Mix (Agilent Technologies, Inc., Santa Clara, CA, USA) as described previously (Ninomiya et al, 2020). Relative expression ratios were calculated using the comparative cycle threshold (Ct) method. The actin-encoding gene was used as a normalization reference. Each sample was tested in triplicate, and the standard deviation is presented. The primer sets used were described in Hagiwara et al, 2017.

### Microsatellite typing

Microsatellite typing was performed as described previously (Hagiwara et al, 2014). Briefly, nine microsatellite regions of approximately 400 bp were PCR amplified using purified genome DNA as a template and sequenced by the Sanger method. The repeat numbers of each locus were counted from the sequences. A dendrogram was constructed using Cluster 3.0 by hierarchical clustering with City-block distance for average linkage, and drawn using Treeview ver. 1.1.6r2 (de Hoon et al, 2004, Saldanha, 2004).

### Genome sequencing

Whole-genome sequencing using next-generation methods was performed as described previously (Hagiwara et al., 2018). In brief, we extracted genomic DNA from overnight-cultured mycelia with NucleoSpin Plant II (Takara Bio, Shiga, Japan). For paired-end library preparation, an NEBNext Ultra DNA Library Prep Kit (New England BioLabs, MA, USA) and NEBNext Multiplex Oligos (New England BioLabs) were used in accordance with the manufacturer’s instructions. A total of 11 strains including 3-1-A to 3-1-H, IFM 59365, IFM 61407, and IFM 61578 were sequenced. Paired-end sequencing (150-bp) on a HiSeq 4000 system (Illumina, San Diego, CA, USA) was carried out by GENEWIZ (South Plainfield, NJ, USA).

### SNP detection

In addition to the abovementioned 11 strains, we used raw data for seven strains for comparison, which have been taken in a study by Takahashi-Nakaguchi et al (2015). Adapters and low-quality bases from Illumina reads were trimmed by fastp (ver. 0.20.1) (Chen et al., 2018). Filtered reads were aligned against the *A. fumigatus* strain Af293 reference genome using BWA (ver. 0.7.17-r1188) (Li and Durbin 2009). SNP detection was performed as described previously (Hagiwara et al., 2018). Briefly, SNPs were identified by using SAMtools (ver. 1.9) (Li et al., 2009) and filtered with >20-fold coverage, >30 mapping quality, and 75% consensus using in-house scripts (Suzuki et al., 2014; Tenaillon et al., 2012).

### Phylogenetic tree construction

Among the strains that were sequenced, mitochondrial genomes of 12 strains were available and aligned by MAFFT (ver. 7.475) (Katoh and Standley, 2013). A phylogenetic tree was constructed using multithreaded RAxML (ver. 8.2.12) (Stamatakis, 2014), the GTRCAT model, and 1,000 bootstrap replicates, and visualized by iTOL (Letunic and Bork 2019). For chromosomal genome phylogenetic classification, 90,987 polymorphic loci were predicted from 18 strains and concatenated, then used for construction of a phylogenetic tree by the methods described above.

### Genome assembly and gene prediction

Mitochondrial genomes were assembled and annotated using GetOrganelle (ver. 1.6.4) (Jin et al., 2020) and MITOS2 (Bernt et al., 2013), respectively. To filter the mitochondrial reads, trimmed reads were aligned against mitochondrial genomes by BWA (ver. 0.7.17-r1188) (Li and Durbin, 2009), and the mapped reads were filtered by SAMtools (ver. 1.9) (Li et al., 2009) and SeqKit (Shen et al., 2016). Contigs were assembled by VelvetOptimiser (ver. 2.2.6) (Zerbino and Birney 2008), followed by generation of a simulated mate-paired library using wgsim (ver. 0.3.1-r13) (https://github.com/lh3/wgsim). The assembly of nuclear genomes was carried out by ALLPATHS-LG (ver. R52488) (Gnerre et al., 2011). The annotation of assembled nuclear genomes was performed by the Funannotate pipeline (ver. 1.7.4) (https://funannotate.readthedocs.io/en/latest/) as described previously (Takahashi et al., 2021). Following identification of repeat sequences by RepeatModeler (ver. 1.0.11) (http://www.repeatmasker.org/RepeatModeler.html) and RepeatMasker (ver. 4.0.7) (https://www.repeatmasker.org), Funannotate *ab initio* prediction was performed with the option “--busco_seed_species=aspergillus_fumigatus” by Augustus (ver. 3.3.3) (Stanke et al., 2006), GeneMark-ES (ver. 4.38) (Ter-Hovhannisyan et al., 2008), GlimmerHMM (ver. 3.0.4) (Majoros et al., 2004), and SNAP (ver. 2006-07-28) (Ian, 2004) using exon hints from the proteins of *A. fumigatus* Af293 and *N. fischeri* NRRL 181 downloaded from the *Aspergillus* Genome Database (http://www.aspgd.org/) (Cerqueira et al., 2014). The completeness of draft genomes and predicted proteins was evaluated by BUSCO (ver. 4.0.6) (Seppey et al., 2019) with the database eurotiales_odb10. Most tools were obtained through Bioconda (Grüning et al., 2018).

### Detection of orthologous genes

Orthologous relationships with *A. fumigatus* Af293 were determined by RBH with criteria BLASTp (ver. 2.9.0+) coverage >80% and identity >80% (Camacho et al., 2009).

### Visualization of genome-wide distribution of SNPs and orthologous genes

SNP frequency in each 1-kb window was calculated and plotted in 250-bp steps using Python (Van Rossum and Drake 2009) and R (R Core Team 2019) scripts. The orthologous genes of *A. fumigatus* Af293 in each strain were visualized by R script.

## Data availability

The genome sequencing data are deposited to DDBJ as DRA011961. BioSample accession(s): SAMD00322244-SAMD00322251.

## Author contributions

HT and DH designed the research; HT, SO, YK, SU, and DH performed experiments; HT contributed new materials/tools; HT and DH analyzed data; and HT and DH wrote the manuscript.

## ACKNOWLEDGMENTS

This study was supported by a grant from the Institute for Fermentation, Osaka (to DH). HT was partly supported by the National Bioscience Database Center (NBDC) of the Japan Science and Technology Agency (JST), and JSPS KAKENHI Grant Numbers 21K07001 and 16H06279. DH and HT were partly supported by AMED, Grant Number JP19fm0208024. We thank Edanz (https://jp.edanz.com/ac) for editing a draft of this manuscript. The authors declare no competing interest.

## Supporting Information

**Figure S1**. Genome-wide SNP frequency compared among the eight strains (3-1-A to 3-1-H). The 10 regions where the pattern is characteristically distinct among the strains are marked by red boxes.

## Notes

### Competing Interest Statement

The authors have declared no competing interest.

