## Supplementary figures and images for "Intimate genetic relationships and fungicide resistance in multiple strains of human pathogenic fungus *Aspergillus fumigatus* isolated from a plant bulb"

### Fig. S1

#### Slide 1
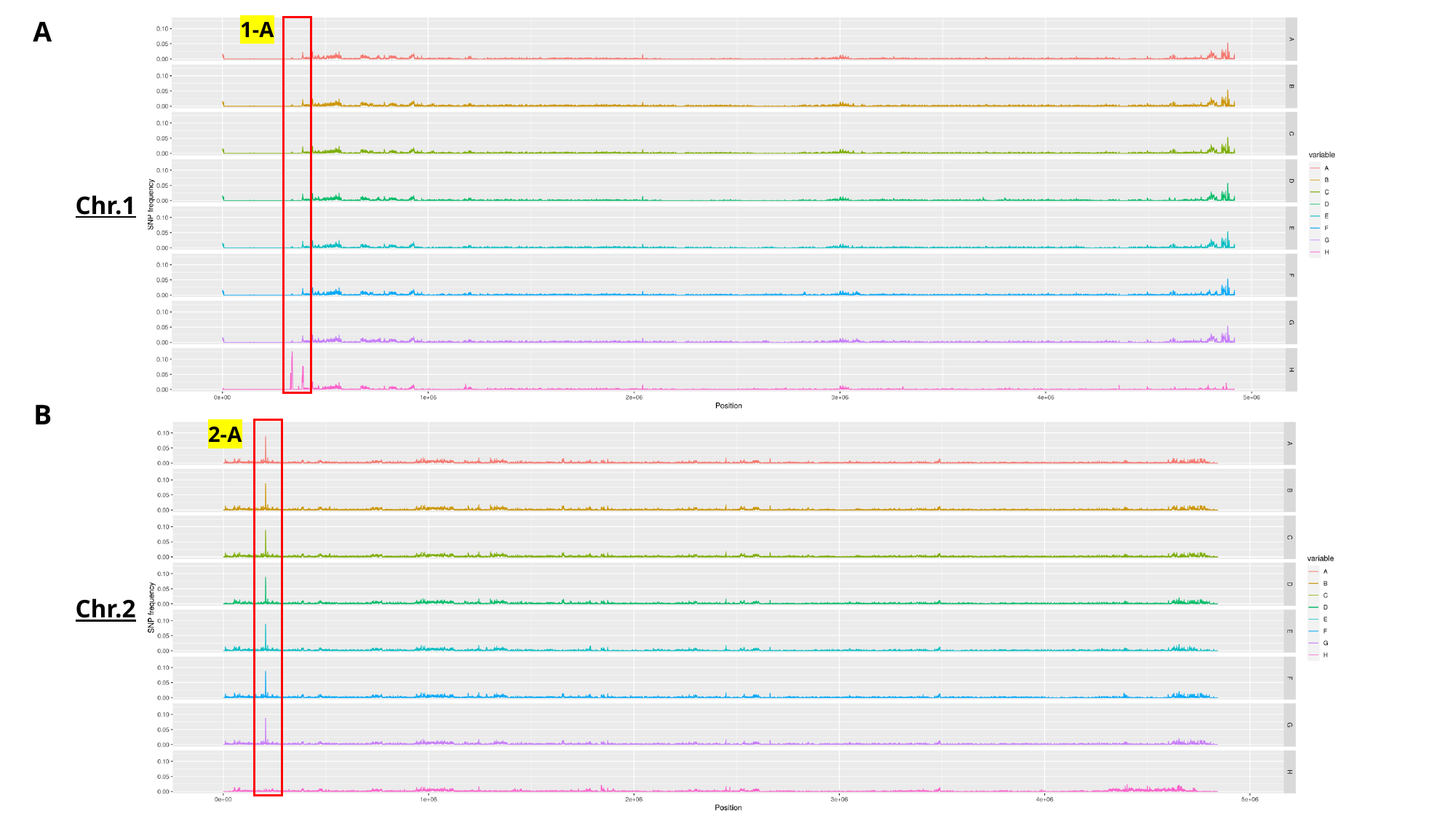

A
1-A
Chr.1
B
2-A
Chr.2

#### Slide 2
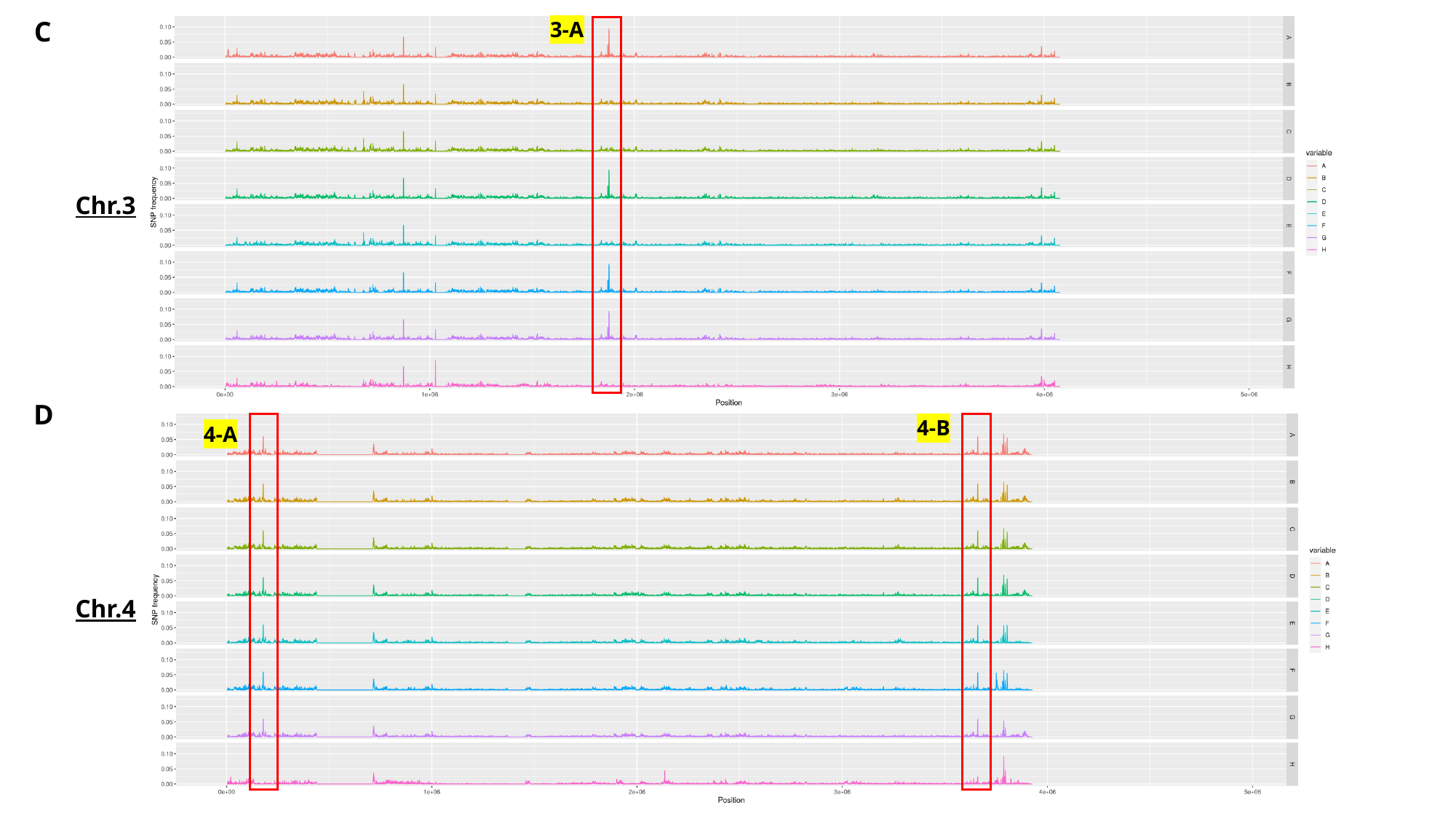

C
3-A
Chr.3
D
4-B
4-A
Chr.4

#### Slide 3
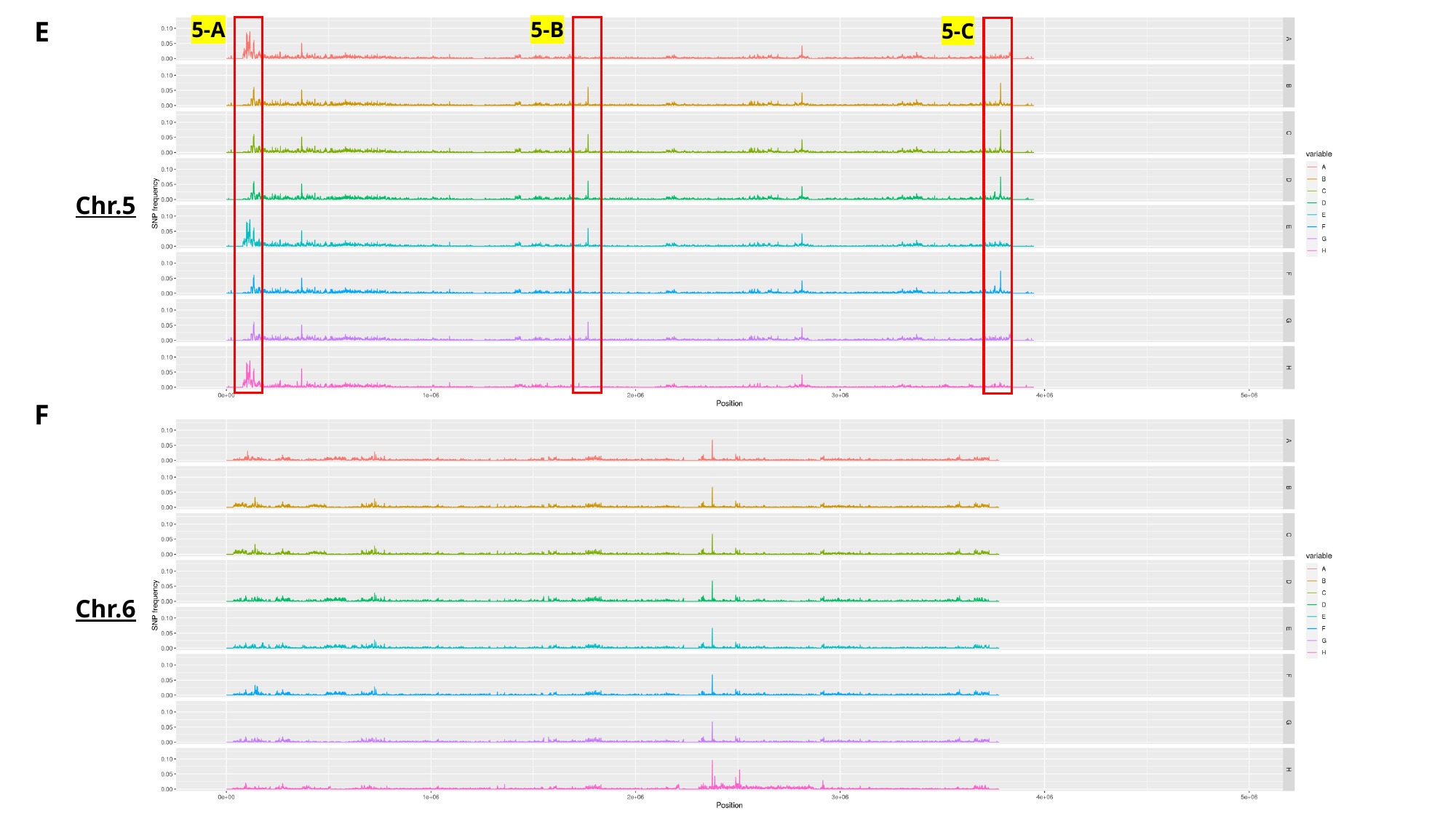

E
5-A
5-B
5-C
Chr.5
F
Chr.6

#### Slide 4
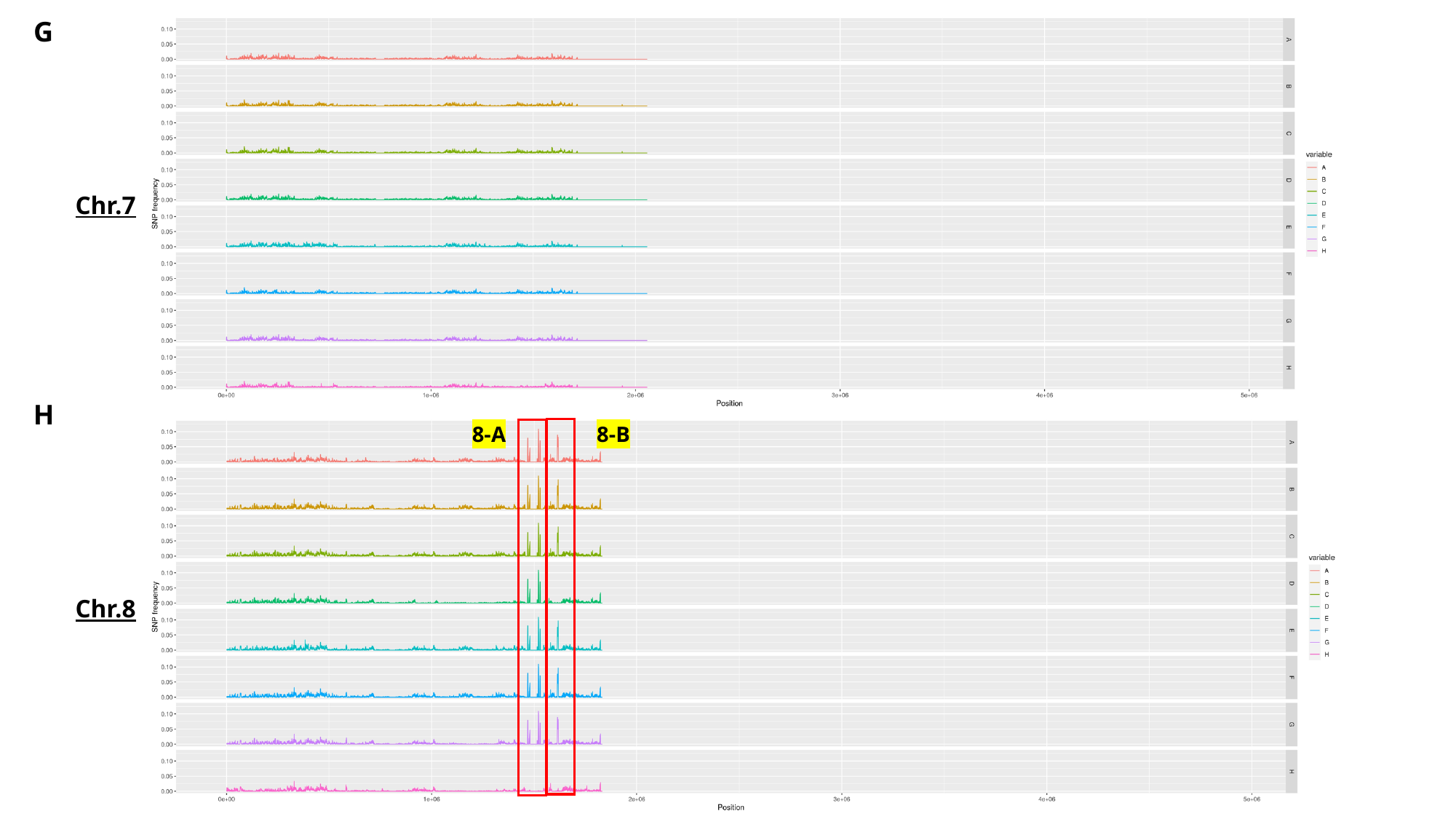

G
Chr.7
H
8-A
8-B
Chr.8
